# Is Vagus Nerve Stimulation Brain Washing?

**DOI:** 10.1101/733410

**Authors:** Kevin P. Cheng, Sarah K. Brodnick, Stephan L. Blanz, Weifeng Zeng, Jack Kegel, Jane A. Pisaniello, Jared P. Ness, Erika Ross, Evan N. Nicolai, Megan L. Settell, James K. Trevathan, Samuel O. Poore, Aaron J. Suminski, Justin C. Williams, Kip A. Ludwig

**Author notes:** Authors contributed equally to this work (Kevin P. Cheng, Sarah K. Brodnick, Justin C. Williams, Kip A. Ludwig).

## Abstract

Vagal nerve stimulation (VNS) is an FDA approved treatment method for intractable epilepsy, treatment resistant depression, cluster headaches and migraine with over 100,000 patients having received vagal nerve implants to date. Moreover, evidence in the literature has led to a growing list of possible clinical indications, with several small clinical trials applying VNS to treat conditions ranging from neurodegenerative diseases to arthritis, anxiety disorders, and obesity. Despite the growing list of therapeutic applications, the fundamental mechanisms by which VNS achieves its beneficial effects are poorly understood and an area of active research. In parallel, the glymphatic and meningeal lymphatic systems have recently been proposed and experimentally validated to explain how the brain maintains a healthy homeostasis without a traditionally defined lymphatic system. In particular, the glymphatic system relates to the interchange of cerebrospinal fluid (CSF) and interstitial fluid (ISF) whose net effect is to wash through the brain parenchyma removing metabolic waste products and misfolded proteins from the interstitium. Of note, clearance is sensitive to adrenergic signaling, and a primary driver of CSF influx into the parenchyma appears to be cerebral arterial pulsations and respiration. As VNS has well-documented effects on cardiovascular and respiratory physiology as well as brain adrenergic signaling, we hypothesized that VNS delivered at clinically derived parameters would increase CSF influx in the brain. To test this hypothesis, we injected a low molecular weight (3 kD) lysine-fixable fluorescent tracer (TxRed) into the CSF system of mice with a cervical vagus nerve cuff implant and measured the amount of CSF penetrance following VNS. We found that the clinical VNS group showed a significant increase in CSF dye penetrance as compared to the naïve control and sham groups. This study demonstrates that VNS therapeutic strategies already being applied in the clinic today may induce intended effects and/or unwanted side effects by altering CSF/ISF exchange in the brain. This may have broad ranging implications in the treatment of various CNS pathologies.

**One Sentence Summary:** Cervical vagus nerve stimulation using clinically derived parameters enhances movement of cerebrospinal fluid into the brain parenchyma presenting a previously unreported effect of vagus nerve stimulation with potential clinical utility.

## INTRODUCTION

Neuromodulation by vagus nerve stimulation (VNS) dates as far back as the late 1800’s when it was observed that mechanical stimulation of the vagal region suppressed seizure activity*(1)*. Throughout the mid-to-late 20^th^ century, studies performing electrical VNS to better understand the functional anatomy of the autonomic nervous system demonstrated VNS can influence brain activity and bloodflow*(2–5)*. Today, VNS is an FDA-approved treatment for refractory epilepsy, treatment resistant depression, cluster headaches and migraine. Further, VNS has been proposed as a potential therapeutic option in an ever-expanding list of conditions including Alzheimer’s Disease, Bipolar Disorder, and obesity*(6)*.

Despite this growing list of applications, the therapeutic mechanisms of VNS are not well understood, in part due to the highly complex and mixed nature of the vagus nerve’s connective domain. The vagus nerve (cranial nerve X) is made up of a mix of efferent (∼20%) and afferent (∼80%) fibers within the parasympathetic and sympathetic peripheral nervous system*(7, 8)*. The vagal trunk gives rise to multiple efferent branches that innervate organs such as the larynx, pharynx, heart, lungs, gastrointestinal tract, various glands and involuntary muscles. Vagal cholinergic efferents exert systemic immunomodulatory effects and constitute an important line of communication in the brain-gut axis*(9, 10)*. Vagal afferents pass through the nucleus tractus solitarius (NTS) to a diverse array of brain structures including locus coeruleus (LC) and the dorsal raphe nuclei through which they influence noradrenergic and serotonergic neurotransmitter levels within the brain*(6)*. Clinical investigations utilizing VNS have demonstrated that it can exert significant cardiovascular and respiratory effects, of which the most commonly reported are bradycardia, dyspnea, and hypotension*(11–13)*.

In parallel, over the past several years the clearance systems of the brain have received increasing attention as a subject of scientific study and as a potential target for therapeutic treatments for Alzheimer’s Disease, Traumatic Brain Injury, and Epilepsy. Historically, the mechanisms by which the brain clears itself of extracellular waste and maintains homeostasis have remained poorly defined, as the brain does not contain the traditionally defined lymphatic vessels that function to maintain tissue homeostasis in the periphery. Recently, however, two novel systems for the clearance of misfolded proteins and cellular waste products from the brain, the glymphatic and meningeal lymphatic systems, have been described*(14, 15)*. The glymphatic clearance system, so-named due to its dependence on astroglial aquaporin-4 (AQP4), describes the interchange of cerebrospinal fluid (CSF) and interstitial fluid (ISF) to clear interstitial space of cellular waste and misfolded proteins*(14)*. These clearance pathways fit within a larger CNS lymphatic system that involves the production of CSF, its movement through the ventricles, subarachnoid space, and into the brain parenchyma wherein the CSF mixes with interstitial fluid collecting solutes from the extracellular milieu and finally draining the CSF/ISF/solute mixture through various routes*(16)*. This collective system is believed to have particular relevance in the area of neurodegenerative diseases, such as Alzheimer’s (AD) and Parkinson’s (PD) where the build-up of misfolded proteins is thought to be a significant contributor to the disease’s progression*(14, 17)*.

The CSF/ISF clearance system is affected by a wide array of lifestyle factors such as sleep, body position, age, and alcohol consumption*(18–21)*. In addition, this system has been shown to become compromised in pathological states following traumatic brain injury (TBI) and in AD progression*(22, 23)*. Thus, it has been suggested that improvement or rescue of clearance function in these pathologies may be of potential therapeutic benefit. Importantly, aspects of the brain clearance systems are sensitive to both changes in noradrenergic signaling associated with sleep/arousal states and to cerebral arterial pulsatility which is hypothesized to drive CSF penetrance into the parenchyma*(18, 24–26)*. Given the well-known noradrenergic, cardiovascular, and respiratory effects induced by VNS, we theorized that VNS may be able to increase CSF penetrance, thereby providing a previously unknown and unexplored therapeutic mechanism of the reported beneficial outcomes of VNS*(4, 12, 27, 28)*. Here, we measured brain-wide changes in CSF penetrance, a component of the glymphatic system, in response to cervical VNS using clinically derived stimulation parameters.

## RESULTS

### VNS increases CSF-penetrance to the brain parenchyma

To understand the impact of clinical VNS paradigms on CSF penetration into the brain parenchyma, the VNS implantation procedure and stimulation parameters were selected to mimic clinical use of VNS as closely as possible. In clinical practice to treat epilepsy/depression there is no interoperative functional mapping to guide electrode placement on the surgical vagus. After implantation the stimulation amplitude is placed at the highest level that is tolerable to the patient, without titrating to optimize changes in heart rate or breathing, with the presumption that the highest tolerable amplitude will activate the largest number of vagus nerve afferent fibers. Clinical VNS to treat epilepsy is typically performed at 30 Hz with pulse-widths most frequently in the range of 100 - 500 µS and a 10% duty cycle intended to minimize side-effects*(29)*. Optimization studies in rodents have demonstrated an inverted U-function in regard to VNS and cortical plasticity with a maximal effective amplitude at 800 uA*(30)*. Thus, we implanted mice with cervical vagus nerve cuffs and applied electrical stimulation at an amplitude of 800 µA with 200 µS pulse-width at 30 Hz with a 10% duty cycle (30 seconds ON, 4 minutes 30 seconds OFF). Additionally, recent reports have indicated that 40 Hz entrainment by stimuli delivered continuously using both optogenetics and external auditory and visual sensory stimuli reduced amyloid plaque deposition and improved cognitive performance in a mouse model of AD*(31, 32)*. To test whether VNS delivered continuously at 40 Hz may achieve similar results we also applied a second stimulation paradigm of 40 Hz continuous VNS (200 µS pulse-width) over 1 hour. In this second cohort, the stimulation amplitude was adjusted to the lowest amplitude at which an observable VNS-induced cardiovascular response was seen. This was done as we found that the 800 uA amplitude used in the clinically derived periodic paradigm was not tolerable when delivered continuously at 40 Hz over 1 hour; which is perhaps not surprising in light of the intimate connections between the vagus, heart, and lungs and that the 800 uA amplitude was derived from studies in which VNS was only applied for a short period of time*(30)*.

Following implantation, we tested engagement of the vagus nerve by the cuff electrode by measuring induction of cardiovascular and respiratory changes in response to stimulation using a pulse oximeter. The stimulation paradigm applied resulted in a range of changes to heart (HR) and respiratory rate (RR) for the duration of the stimulus followed by a rapid recovery once stimulation was stopped (Figure 1C, Supplemental Figure 2). Additionally, visual confirmation of the Hering-Breuer reflex, which causes a brief cessation of breathing to prevent lung over-inflation, and is characteristic of VNS, was observed in most but not all animals. After confirming engagement, experimental animals received the clinically derived VNS over the course of 1 hour, and the degree of CSF penetrance was quantified by the spread of a fluorescent dye injected into the cisterna magna (CM) using previously established methods*(14, 33)*. Following perfusion, the fractional area to which the dye had spread was measured using a thresholding algorithm as previously described*(14)*. A total of 12 – 16 100um thick slices corresponding to the region spanning +1 to −2 mm relative to bregma were analyzed for each animal. We found that the clinically derived periodic VNS parameters significantly increased the degree of CSF dye penetrance (18.60% ± 1.98%) relative to naïve controls (10.75% ± 1.26%, p < 0.001), the sham VNS (13.05% ± 2.29%, p = 0.006), and the continuously delivered 40 Hz VNS (12.78% ± 1.64%, p = 0.046) (Figure 1A,B). Whereas there was no difference between the naïve controls, sham VNS (p = 0.997), and continuous 40 Hz VNS (p = 0.967) groups.

**Figure 1:**
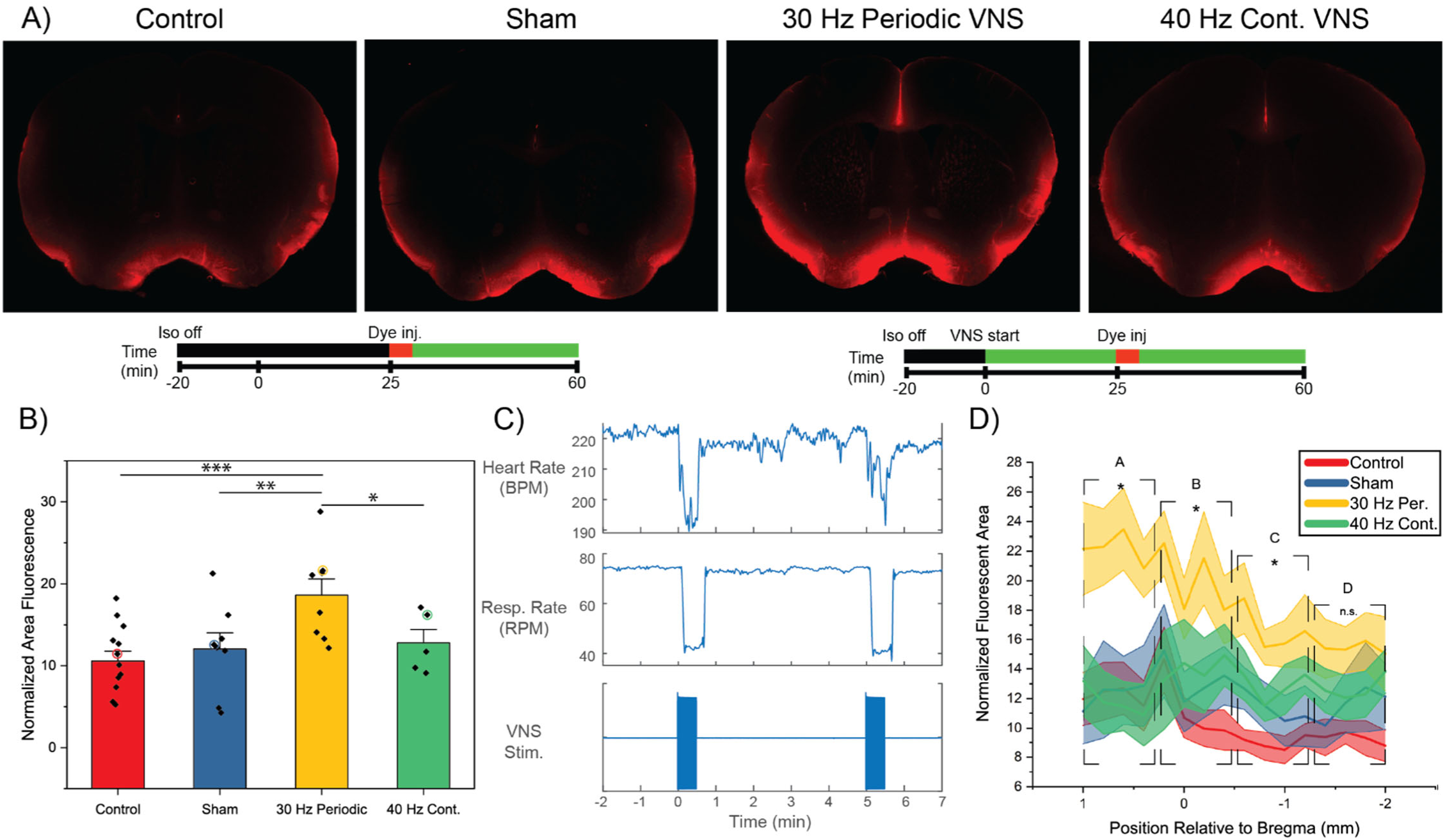
Vagus nerve stimulation increases CSF dye penetrance. A) Representative images of whole slice images showing difference in CSF penetrance in response to VNS as compared to control and sham. Stimulation parameters were as follows: Biphasic, 30 Hz, amplitude = 800 µA, pulse width = 200 uSec with 30 seconds ON cycled every 5 minutes. The fluorescent portion of each slice was quantified as a function of the total slice area. A total of 12-16 slices from between the regions of 1 and −2 mm relative to bregma were assessed per animal in this way. Colored bar beneath shows the relative timeline for each group. In order to account for time of anesthesia a waiting period was incorporated into the control and sham groups. B) Average degree of CSF penetrance across groups, dots represent individual animals in each group. Colored circles correspond to the animals from which the slice images in (A) were taken. C) Representative traces of heart and respiratory rates in response to VNS. D) VNS increases CSF penetrance more in anterior regions of the brain than posterior. Positional slice-by-slice representation of the normalized fluorescent area at each position relative to bregma. Solid line represents the average normalized fluorescent area of all animals at that section per condition (shaded area = ±1 S.E.M). For analysis the data was subdivided into 4 equivalently sized bins which were individually analyzed. Split-plot design linear mixed model analysis with Sidak post-hoc correction. Error bars = ±1 S.E.M. Control n = 12, sham n = 8, 30 Hz periodic VNS n = 8, 40 Hz continuous VNS n = 5, *** p < 0.001, ** p < 0.01, * p < 0.05

Further, measurement of CSF-dye penetrance at each slice position relative to bregma, rather than as an aggregate per animal, revealed that the VNS induced increase in CSF penetrance was more prominent in the anterior portion of the region analyzed. Sections from each animal were combined into four equivalently sized positional bins and analyzed across experimental groups. We found that VNS induced significant increases in CSF penetrance in the three anterior bins but not in the fourth, most posterior, bin (Figure 1D). The more pronounced VNS-induced CSF penetrance found in the anterior parts of the brain may be due to their closer proximity to the middle cerebral artery (MCA) and branching arterioles along which CSF is hypothesized to travel in the glymphatic model*(34)*. This may suggest that the mechanism of increased CSF penetrance is tied to VNS cerebral hemodynamic effects. Supporting this idea, vagal connections to the sphenopalatine ganglion (SPG), which directly parasympathetically innervates the anterior circulation, are well-documented*(35, 36)*. However, similar anterior/posterior distributions have been observed in other studies where VNS was not present. Consequently, anterior/posterior distribution of increased CSF penetration may simply be an artifact of the site of injection and relative flow direction of CSF in an anterior to posterior direction*(37, 38)*.

### VNS-induced CSF penetrance could not be directly linked to cardiac and respiratory responses

In order to better understand the mechanism behind VNS-induced CSF penetrance, we attempted to correlate the degree of CSF penetrance with the physiological responses recorded during stimulation. The CSF influx hypothesized to occur in the glymphatic model is driven by cerebral arterial pulsation and it was recently reported that increased glymphatic influx was correlated with a reduced HR in mice under anesthesia*(24, 26, 38)*. Although no direct experiments have been undertaken to assess the effect of VNS on cerebral arterial pulsation, several studies detail the systemic cardiovascular and respiratory responses of VNS*(12, 13)*. In the process of testing vagus nerve engagement our results also demonstrated the well-documented cardiovascular and respiratory effects of VNS. While the responses were typically a suppression of both HR and RR during stimulation there was wide variability in the magnitude of VNS effects across animals (Table 1) and no obvious correlation could be found with any specific physiological response and CSF penetrance (Supplemental Figure 3). This variability may have arisen from differences in electrode-nerve contact, the cuff’s position along the vagal trunk during implantation, or in the anatomical layout of baroreceptor fibers and the aortic depressor nerve (ADN) between animals. Indeed, the inconsistency of physiological responses across the VNS group are reflective of previous studies which have shown large inter-subject variability in the response to VNS across a range of animal models and in humans*(39–41)*. In pigs and dogs, the ADN at the cervical level may run with or independently of the vagal trunk*(39, 40)*. As the intent of this study was to mimic clinical VNS for its FDA-approved indications, electrode location and stimulation parameters were deliberately not optimized to isolate and/or maximize changes in heart rate or changes in breathing rate to match clinical practice*(29)*. These inter-subject variances may explain why that, although the aggregate data across groups demonstrated significant increases in CSF penetrance due to VNS, taken individually there was a wide range in the degree of CSF penetrance within the clinical VNS group with some animals exhibiting similar levels as the control and sham groups (Figure 1B).

**Table 1:**
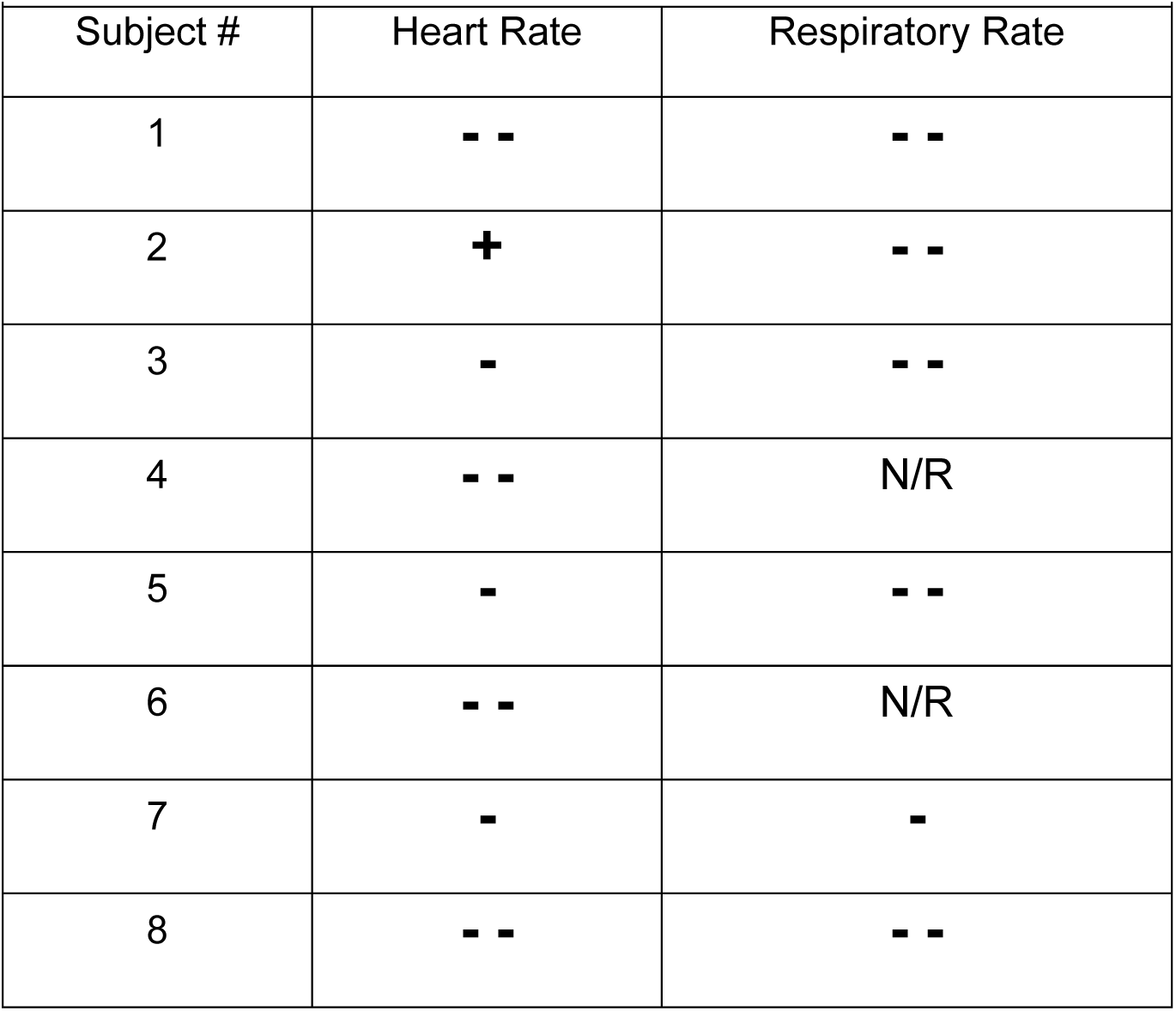
Individual systemic cardiopulmonary responses to VNS. Individual changes to heart rate (HR) and respiratory rate (RR) observed in response to VNS. Measurements were recorded using a pulse-oximeter with the sensor placed on the thigh. Single symbols denote changes of < 10% of baseline, double symbols represent changes > 10% baseline, N/R denotes responses during which the pulse oximeter was not able to get an accurate reading.

| Table 1: Individual systemic cardiopulmonary responses to VNS |  |  |
| --- | --- | --- |
| Subject # | Heart Rate | Respiratory Rate |
| 1 | - - | - - |
| 2 | + | - - |
| 3 | - | - - |
| 4 | - - | N/R |
| 5 | - | - - |
| 6 | - - | N/R |
| 7 | - | - |
| 8 | - - | - - |

## DISCUSSION

The brain’s waste clearance systems are increasingly appreciated as a critical factor in the maintenance of healthy brain homeostasis and conversely in the progression of neuropathological processes (i.e. the build-up of misfolded proteins in neurodegenerative diseases). In parallel, neuromodulation of the vagus nerve has been demonstrated to be clinically effective across a large and continuously growing number of indications yet the mechanisms behind these beneficial outcomes remain poorly understood. Here, our results demonstrate for the first time that VNS using clinically derived parameters can alter penetrance of CSF into the brain parenchyma and may thus provide a novel method for increasing CSF/ISF exchange and brain waste clearance.

In our retrospective analysis VNS-induced CSF penetrance could not be correlated with a specific effect of VNS on HR and RR, however due to the limited scope of this study such a relationship cannot be ruled out. Here, we primarily used the physiological responses to confirm engagement of the vagus nerve following cuff implantation, but future prospective studies should be performed isolating and optimizing each physiological response to VNS with a sufficient number of animals to achieve appropriate statistical power. A given subject’s response to VNS can vary widely as a result of individual anatomy, cuff placement along the vagal trunk, and electrode-nerve contact, and to better tease apart the potential relationship between VNS-induced physiological responses these factors must be tightly controlled. While cuff placement and electrode-nerve contact may be better controlled via surgical and cuff design methods, addressing inter-subject anatomical variability is inherently more difficult. Since it is impossible to know an individual subject’s response to VNS *a priori*, this would likely require indeterminately expanding the number of experimental subjects to sufficiently populate experimental groups with different physiological responses (i.e. VNS-induced HR increase vs. HR decrease) for statistical analyses.

The VNS-induced effects on CSF penetrance may also be partially or completely independent of VNS systemic cardio and respiratory responses and depend on more regionalized brain specific VNS effects. The wide number of CNS targets connected to the vagus nerve and the expanding number of factors known to influence the brain’s waste clearance systems certainly leaves open the possibility of an alternative mechanism. For example, transient changes in cerebral arterial blood flow due to VNS, as they are mediated in part through cross connectivity to the facial nerve and subsequent parasympathetic innervation of the cerebral artery, would not be expected to depend on systemic heart rate or breathing responses*(4, 35, 36, 42)*. VNS can also alter the neurotransmitter and metabolite content of CSF possibly leading to unknown interactions with the astrocytic end-feet lining the paravascular space (PVS) that are hypothesized to be the primary cellular regulators of CSF entrance to the brain parenchyma*(14, 43)*. In rats, VNS reduced cortical edema following traumatic brain injury and thereby improved functional recovery*(44)*. Although the exact mechanism was unidentified, it was hypothesized that VNS-induced norepinephrine release from locus coeruleus led to activation of noradrenergic receptors to alter the water and ionic balance of the brain*(44)*. The glymphatic system is heavily influenced by brain noradrenergic signaling although reports indicate that inhibition of noradrenergic signaling leads to increased glymphatic flux*(18, 45)*. VNS has also been shown to acutely decrease intracranial pressure in pigs through an unknown mechanism that was determined to be independent of VNS-induced cardiovascular effects*(46)*. Exploring the multiple pathways, physiological responses to VNS, and their contributions to CSF penetrance will be the goal of future work.

Recent reports indicate that gamma (40 Hz) entrainment is effective at reducing amyloid deposition and improving cognitive performance in AD mouse models through an as yet unidentified mechanism *(31, 32)*. Interestingly, in these studies the beneficial effect was achieved not by direct electrical stimulation but by audial and visual sensory stimuli. In light of these reports and the aforementioned links between brain waste clearance and AD we tested whether VNS delivered continuously at 40 Hz affects CSF penetrance. Our results indicate that, unlike the clinically derived periodic paradigm, continuous 40 Hz VNS was unable to induce increases in CSF penetrance. If the primary mechanism of VNS-induced CSF penetrance is through alterations in cerebral hemodynamics it is possible that this mechanism requires the periodic ‘shock’ to the system leading to large repeated fluctuations in cerebral blood flow whereas a continuously delivered stimuli leading to a sustained alteration of cerebral blood flow may not achieve the same swings in arterial pulsation hypothesized to drive CSF influx. However, as noted, the stimulation amplitudes applied to the continuous 40 Hz group (100 – 300 µA) was not the same as the clinically derived 30 Hz periodic group (800 uA) due to that stimulation amplitude being intolerable to the animal over the course of the experiment. As the parameters for the continuous 40 Hz group were determined by finding the minimum stimulation amplitude to induce a cardiovascular response it is possible that the amplitudes chosen were simply insufficient to activate the vagal pathways involved to increase CSF penetrance. While the cardiovascular response observed indicates at least activation of parasympathetic efferents to the heart, both the degree and cortical sites of VNS-induced alterations in cerebral blood flow have been shown to be amplitude dependent in humans*(47)*. Alternatively, as we only assessed CSF penetrance or the ‘start’ point of CSF/ISF exchange it is possible that 40 Hz gamma entrainment affects a component of the brain waste clearance system at another level which would not have been detected by our methods.

It is important to note that in the present study all experiments were performed under anesthesia considering the proposed importance of arousal state on glymphatic function*(18, 48)*. Further, there have been a number of conflicting reports as to the effect of anesthesia and different anesthetics on CSF influx and efflux from the brain parenchyma*(18, 26, 49, 50)*. Of note, Gakuba et al. found that anesthesia with either ketamine, ketamine/xylazine, or isoflurane significantly inhibits CSF movement as compared to the awake state*(49)*. In contrast, Xie et al. reported that glymphatic function, as quantified by penetrance of a CSF tracer to the parenchyma, was significantly higher in sleep and ketamine/xylazine anesthetized states versus the awake state *(18)*. Recently, Hablitz et al. studied the effect of a range of anesthetic regiments on glymphatic function and found that CSF tracer penetrance was highest under ketamine/xylazine and significantly impaired under isoflurane*(26)*. This may explain the results of Gakuba et al. who performed all tracer injections under isoflurane before switching to their experimental target anesthetic*(26, 49)*. In our study, although surgery was performed under isoflurane, all injections were done under ketamine/dexmedetomidine anesthesia after allowing the isoflurane to wear off.

Separately, Ma et al. reported that in the awake state tracers injected into the CSF are rapidly cleared to the systemic blood via the lymphatic system and that this efflux was greatly impaired under ketamine/medetomidine anesthesia*(50)*. They also observed a negative correlation between tracer found in the systemic blood and within the brain parenchyma. They therefore suggest that the previously reported increases in glymphatic function under sleep and anesthetic states reported by others may have actually been due to the build-up of tracer within the CSF system as a result of the significantly diminished outflow. The overall contribution of the glymphatic system to brain clearance remain a current topic of debate within the scientific community. Our results demonstrate that VNS can alter, at least in an anesthetized state, the penetrance of CSF into the brain parenchyma which has many potential implications outside of the clearance of misfolded proteins. Increasing the amount of CSF in the brain parenchyma presumably would increase the distance between neighboring electrically excitable cells. This would serve to decrease non-synaptic coupling between neurons, also known as ephaptic coupling; ephaptic coupling has been demonstrated to strongly entrain action potentials within proximally located neurons and has been increasingly linked to epileptic pathologies*(51–54)*. Moreover, increasing CSF penetration into the parenchyma presumably alters extracellular concentrations of endogenous neurochemical transmitters and other molecules. Changes in tonic and/or phasic concentrations of neurochemical transmitters and other molecules that modulate synaptic transmission may have profound implications on circuit function, by altering intrinsic excitability within the brain.

Here, only a single, clinically derived, stimulation paradigm was applied despite multiple examples within neuromodulation demonstrating the importance of optimizing the stimulation parameters to achieve the maximal desired effect and avoid noxious, harmful stimuli. It is likely then that the application of VNS to increase brain waste clearance would also benefit from such optimization and future studies should take into consideration testing of multiple frequencies, amplitudes, and duty cycles. Similarly, while we targeted the vagus nerve due to its widespread and growing list of clinical indications it should be noted that other cranial nerve targets have recently come to attention as potential targets for neuromodulation. If the primary mechanism driving CSF entrance into the brain parenchyma is proven to be cerebral arterial pulsation, then the facial nerve (CN VII) becomes a tempting target as it has been well-documented to regulate cerebral arterial dilation and blood flow *(55)*. Interestingly, transection of the facial nerve blocked the ability of VNS to induce cerebral bloodflow indicating the effects of VNS may be due to crosstalk of the vagus and facial nerves since the facial nerve directly parasympathetically innervates the cerebral arteries*(4, 35)*. In addition, the trigeminal nerve (CN V) has similar anatomical connections to key brain nuclei as the vagus, including the NTS and locus coeruleus, and has gained an increased research focus as an alternative to VNS as, in humans, it can be more effectively stimulated by superficial, non-invasive, methods. Recently, trigeminal nerve stimulation was also used to reduce cerebral edema, enhance cerebral bloodflow, and improve recovery following severe TBI in rats*(56)*.

## CONCLUSIONS

In the present study, our results demonstrate that electrical stimulation of the vagus nerve can be used to modulate one aspect of the brain’s waste clearance systems, which are beginning to emerge as critical to the healthy brain’s homeostasis, thus highlighting the need for future studies to both better define the stimulation parameter space and the mechanisms involved. Indeed, the discovery that VNS mimicking clinical use for epilepsy and depression can increase the penetration of CSF into the brain parenchyma has profound implications for the underlying mechanisms for VNS therapies as well as unwanted side effects. At a minimum, the new finding that VNS increases CSF penetration into the parenchyma is a potential confound to studies intended to isolate the circuit pathways altered during VNS to understand mechanism of therapeutic action that must be addressed going forward. As a previously unexplored facet of cranial nerve stimulation, the modulation of CSF penetration could also conceivably lead to unforeseen and unwanted consequences during chronic administration. As such, this phenomenon warrants scientifically rigorous study to isolate the fundamental mechanism causing increased CSF penetration in future studies.

## METHODS

### Study Design

The objective of this study was to determine whether VNS alters CSF penetrance to the parenchyma as a measure of brain waste clearance. To achieve this, we implanted animals with a left cervical vagus nerve cuff and injected a fluorescent tracer into the CSF system at the site of the cisterna magna using previously established methods for measurement of CSF penetrance and glymphatic system function*(14)*. We applied VNS utilizing a clinically derived paradigm to reflect present-day clinical practices as well as in a continuous 40 Hz experimentally derived paradigm.

Sample size was determined based on previous experience in our lab with animal studies. To account for gender and age effects all mice used were male and between the ages of 8 – 14 weeks old. Mice were randomly assigned to experimental groups and controls, shams, and VNS procedures were performed in an interleaved manner. Researchers were not blinded to experimental groups. Animals were excluded from analysis if leakage occurred at the site of injection as indicated by observation of dye fluid external to the atlanto-occipital membrane. Slices were excluded from analysis if they had misfolded during the slide mounting process making area normalization impossible.

### Animal Use

All animal procedures were approved by the Institutional Animal Care and Use Committee (IACUC) at the University of Wisconsin - Madison. All efforts were made to minimize animal discomfort. 8 – 14 week-old C57/BL6J male mice with 12 hour light/dark cycles with ad libitum access to food and water were used. The studies conducted were comprised of four animal groups including a naïve control (n = 12), sham VNS control (n = 8), clinically derived VNS (n = 8), and experimentally derived 40 Hz continuous VNS (n = 5) groups. Control mice underwent the same cisterna magna dye injection procedure and dye injection timeline but did not have a vagus nerve cuff implant. Sham VNS mice received the vagus nerve cuff implant, but the leads were left un-connected and no stimulation was applied.

### Electrical Stimulation of the Vagus

Stimulation of the vagus nerve was applied through a cuff-style electrode made in-house with the body of the cuff made of 500 um inner diameter silicone tubing (AM Systems). Each cuff was made ≈3 mm long and contained two insulated platinum wires (AM Systems) spaced ≈1 mm apart with the insulation within the cuff carefully removed. Stimulation was provided by an RZ2 16-channel stimulator (Tucker-Davis Technologies). Stimulation parameters of the clinically derived group consisted of 200 µs biphasic pulses with an amplitude of 800 µA delivered at 30 Hz and applied at a 10% duty cycle (30 seconds ON and 4.5 minutes OFF) over the course of one hour*(29)*. In clinical practice, the 10% duty cycle is often implemented to minimize side effects while the maximum tolerable amplitude is applied under the assumption that a higher amplitude will maximize engagement and enhance the beneficial effects of VNS. However, optimization studies in rats have demonstrated an inverted U-function in the relationship between stimulation intensity and cortical plasticity with medium range amplitudes (400 – 800 µA) exhibiting optimal effects as compared to low (< 400 µA) and high (> 1.2 mA) amplitudes (Borland 2016). Thus, we chose to apply stimulation at 800 µA to reflect this maximally effective amplitude in rodents while remaining within the range of clinically applied parameters. In the second VNS group stimulation was applied at 40 Hz continuously over the course of one hour to reflect recent studies indicator that gamma (40 Hz) entrainment through various stimuli affect amyloid build-up in a mouse model of AD*(31, 32)*. We found that application of 40 Hz continuous VNS at 800 µA amplitude was intolerable over the duration of the experiment. Thus, stimulation amplitude of this group varied per animal and was adjusted to the minimum amplitude (amplitudes ranged from 100 - 300 µA) at which a cardiovascular response was detected.

Proper nerve engagement was confirmed by monitoring of heart and respiratory rates using a pulse oximeter with an attached thigh sensor (Starr Life Sciences). Pulse oximeter output was plotted and visualized using MATLAB. The output streams of the pulse oximeter corresponding to heart and respiratory rate were aligned with each stimulus event’s onset/offset and plotted against time.

### Cuff electrode implantation procedure

Mice in the two VNS and sham groups underwent a vagal nerve cuff implantation procedure prior to cisterna magna injection. Animals were induced at 5% isoflurane in O_2_ and maintained at 1.75-2.5% isoflurane throughout the surgical cuff implantation procedure. The clavicles, suprasternal fossa, and mid-line of the neck were marked prior to incision. Following mid-line incision, the skin, fatty tissue, and two submandibular glands were separated to reveal the sternohyoideus and sternocleidomastoid muscles. The sternocleidomastoid muscles were then separated from the sternohyoideus to expose the lower edge of the muscle, and just underneath, the carotid sheath containing the vagus. While the internal jugular and carotid arteries were carefully separated no attempt was made to identify and separate the aortic depressor nerve from the vagus to reflect current clinical practice. To maintain contact with the nerve during subsequent manipulations the ends of the cuff were secured with 9-0 suture (Ethicon Inc). Following cuff placement, the area was closed-up while allowing the wires of the cuff to exit externally through the skin.

### Cisterna magna (CM) dye-injection

Injection of a fluorescent tracer into the CSF system via the CM was performed as previously described *(14, 33)*. Briefly, after vagus nerve cuff implantation, the mice were carefully placed into a stereotax with the head at a slight downward angle to provide access to the CM. A straight vertical incision approximately 10mm long was made into the skin just caudal to the occiput and terminating at the atlas. The underlying fascia and muscle tissue was bluntly dissected to expose the CM. An injection port was created using a 30 ga dental needle (Exel International) attached to short length of PE-10 tubing (Instech Laboratories Inc). Using a motorized micro-manipulator (Siskiyou Corp.) the injection port was positioned just posterior to the cisterna magna at a 30-degree angle to the membrane. To prevent leakage of CSF and dye the insertion site was sealed by a small amount of cyanoacrylate glue. The injection port’s tubing was then connected to a syringe pump (Kent Scientific) for dye injection.

Previous reports indicate that the glymphatic system is sensitive to the type of anesthesia applied with isoflurane diminishing CSF penetrance while a combination of ketamine and xylazine was found to be most permissive*(57)*. Further, isoflurane can inhibit synaptic transmission at post-synaptic cleft and is ill-suited for experiments involving electrical stimulation. Thus, following surgery we switched anesthetics from isoflurane to a combination of ketamine and dexmedetomidine (same class of anesthetics as xylazine). After injection of ket/dex, isoflurane was turned off and a delay period was implemented to allow for the effects of isoflurane to dissipate. Care was taken to keep the duration of the delay period across animals consistent. However, due to the differences in complexity of the surgical procedures performed across the experimental groups as well as the time taken to test vagus engagement the delay period duration varied across animals. The variations in delay period between animals was not found to have any effect on CSF penetrance (Supplemental Figure 4). After the delay period, a total of 10 uL of 0.5% TxRed-3kD dextran (Life Sciences) in aCSF was injected into the CM over a 5-minute period and the tracer was allowed to spread for a total of 30 minutes post-injection.

### *Ex vivo* slice preparation and imaging

At the end of the experiment the animals were immediately perfused with saline followed by 4% paraformaldehyde (PFA). The brain was extracted from the cranium and placed into 4% PFA for an additional 24-hour period before being transferred to 1X phosphate buffered saline (PBS). For sectioning, the brains were sliced on a vibratome (Leica Biosystems) into 100 um thick coronal sections spanning the region corresponding to +1 to −2 mm relative to bregma. Starting from the first slice, every other slice was mounted to glass slides with ProLong Gold containing DAPI (Life Sciences), producing approximately 16 analyzable slices per animal. Whole slice images were taken using a fluorescent stereoscope (Nikon SMZ18) with DS-Qi2 camera. All microscope settings including gain and exposure were kept constant across all animals and experimental groups.

### Data and Statistical Analysis

Image data was analyzed using the FIJI version of ImageJ and a thresholding approach as previously described*(14)*. Briefly, regions of interest (ROI’s) encompassing the entire slice were automatically generated by visualization of the DAPI-channel. The ROI’s were then overlaid onto the TxRed-channel images and the percentage of pixels within the ROI above a set threshold was calculated. The threshold was determined by imaging of unstained sections from animals who did not receive the CSF dye and was kept constant across all analyses. A total of 12-16 slices were analyzed per animal, the variability in number was due to tissue sections misfolding during the mounting procedure and excluded from analysis.

To assess global effects of VNS on CSF penetrance the fractional positive fluorescent area of all slices from each group were assessed using a split-plot design linear mixed model with an AR(1) covariance matrix and subsequent pairwise comparisons with Sidak correction for multiple comparisons performed in SPSS (IBM). For the anterior-posterior positional analysis, the same methods were used but the slices were organized into four equivalently sized non-overlapping bins spanning the entire region. Results were considered statistically significant with p < 0.05.

