## SupplementalMaterials for "Is Vagus Nerve Stimulation Brain Washing?"

**Supplemental Materials**

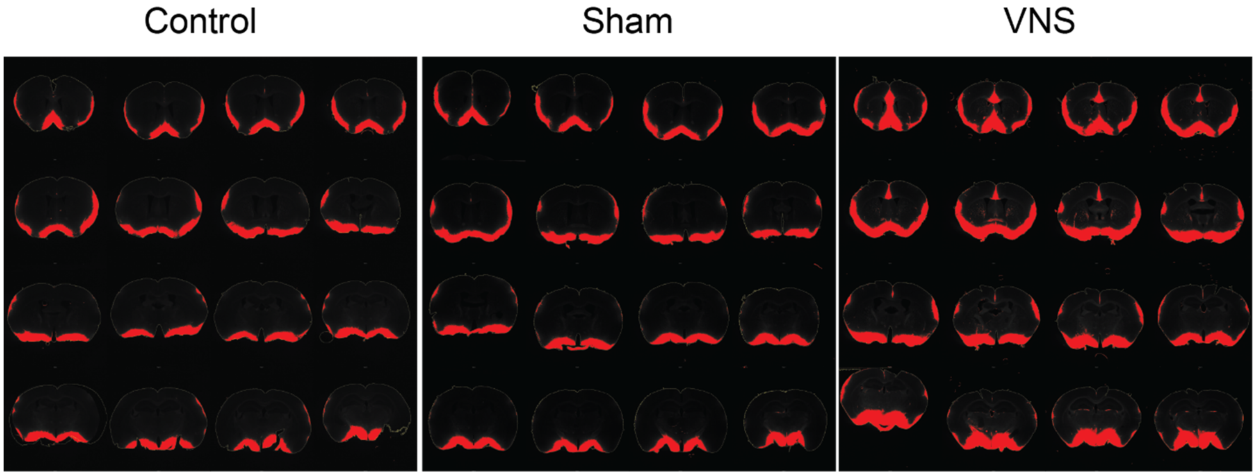


Supplemental Figure 1: Results of thresholding algorithm applied to the full set of slices from animals from each experimental group. Slices were 100 um thick and collected every 200 um from the region spanning -2 to 1 mm relative to bregma. Red filled areas indicate brain regions positive for the CSF tracer as determined by the thresholding algorithm. The threshold was determined by imaging of sections from animals in which no tracer was injected.

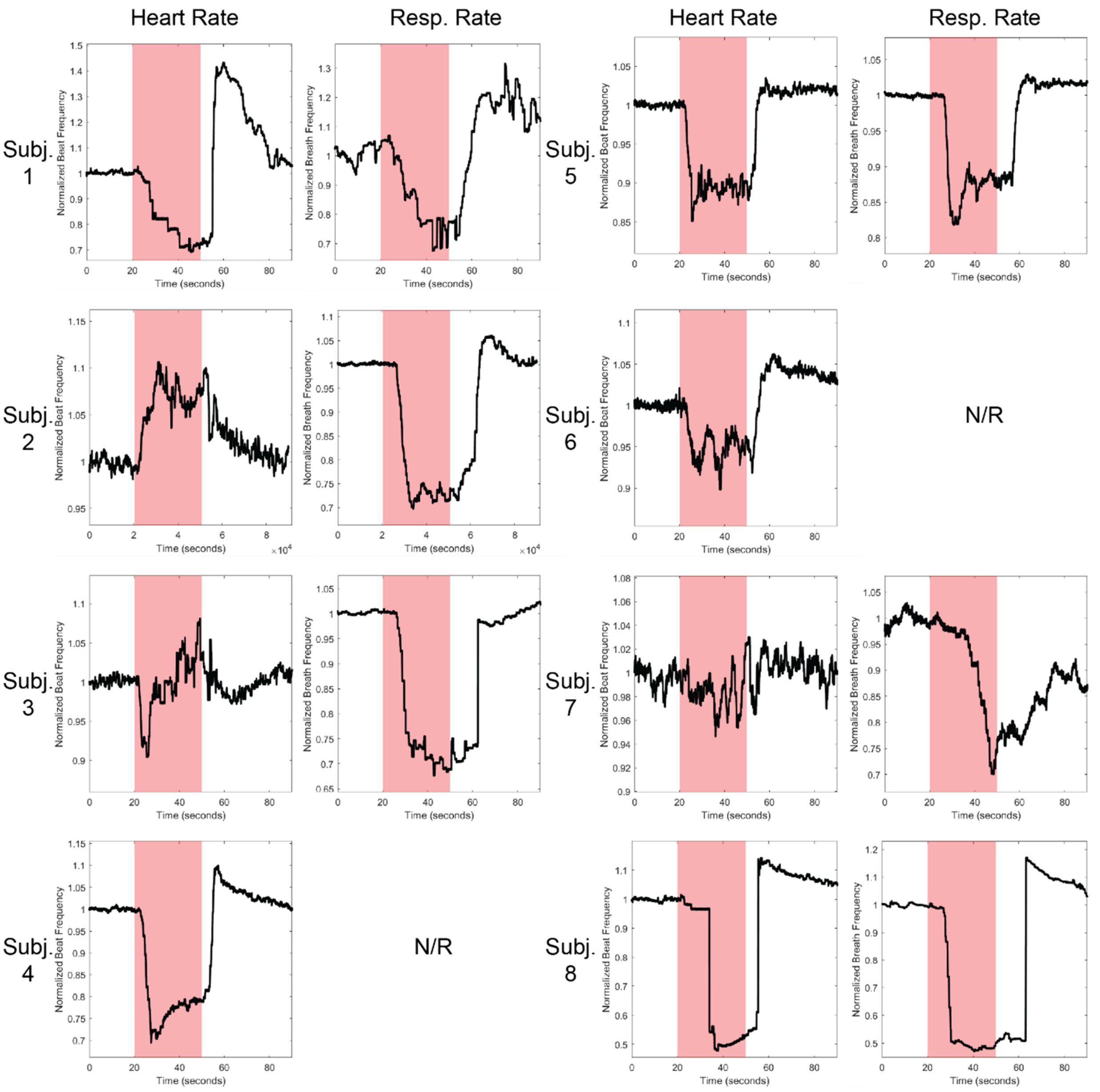


Supplemental Figure 2: Individual heart (HR) and respiratory (RR) rate responses to VNS. Changes in HR/RR for each animal were normalized to baseline and averaged across multiple stimulation events. Areas shaded in red indicate the period of VNS. N/R = not recorded due to equipment error.

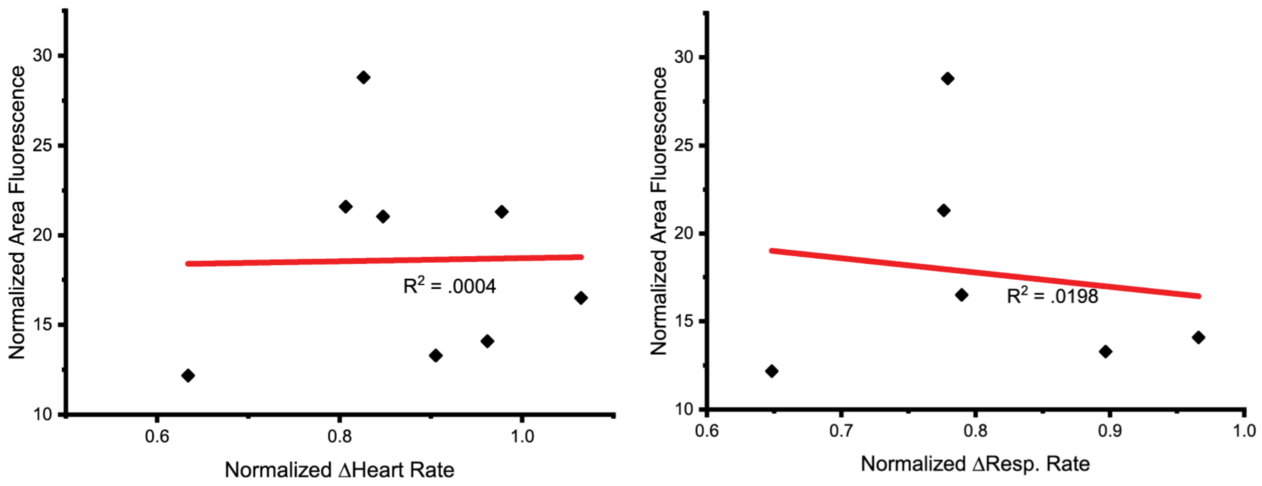


Supplemental Figure 3: Individual measurements of cardiopulmonary responses to VNS was plotted against the degree of CSF penetrance and a linear regression analysis was performed. We found no correlation between the magnitude of the cardiopulmonary response to VNS with the degree of CSF penetrance. However, it should be noted that HR/RR measurements were taken primarily as a means of confirming engagement with the vagus and this study was not designed to rigorously test the relationship between CSF penetrance and VNS effects on heart and respiratory rate.

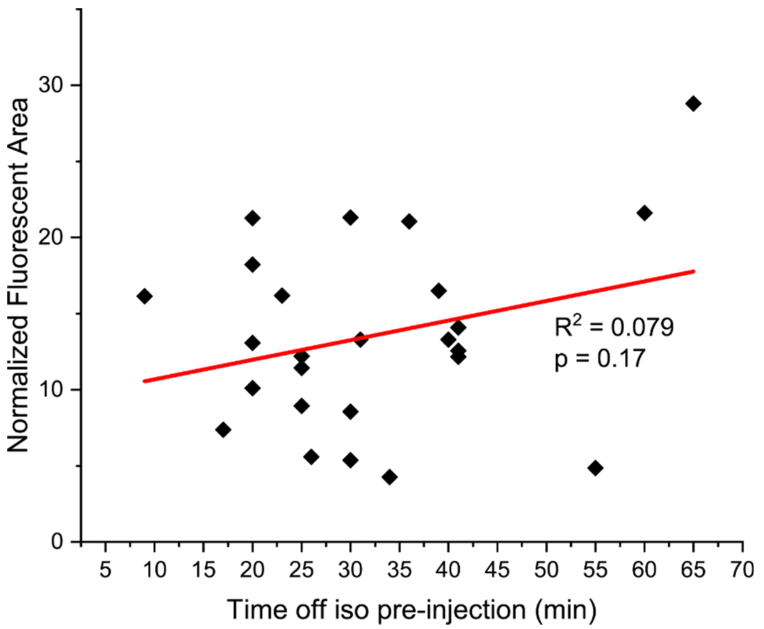


Supplemental Figure 4: The duration of the delay period, corresponding to the time off of isoflurane prior to injection of the CSF tracer, did not have an effect on the degree of CSF penetrance. To account for the experimental variance in the time off of isoflurane across animals the individual delay period and normalized fluorescent area from animals across all experimental groups was combined and plotted. Linear regression analysis revealed no relationship between the time off of isoflurane and the degree of CSF penetrance.
